# An integrated analysis of the epigenetic, genetic, and transcriptional patterns associated with outcome across cancer types

**DOI:** 10.1101/186528

**Authors:** Joan C. Smith, Jason M. Sheltzer

## Abstract

Successful treatment decisions in cancer depend on the accurate assessment of patient risk. To improve our understanding of the molecular alterations that underlie deadly malignancies, we analyzed genomic profiles from 33,036 solid tumors with known patient outcomes. Contrary to expectations, we find that mutations in cancer driver genes are almost never associated with patient survival time. In contrast, copy number changes in these same genes are broadly prognostic. Analysis of methylation, microRNA, mRNA, and protein expression patterns in primary tumors define several additional prognostic patterns, including signatures of tumor mitotic activity and tissue de-differentiation. Co-expression analysis with a cell cycle meta-gene distinguished proliferation-dependent and ‐independent prognostic features, allowing us to construct multivariate survival models with improved stratification power. In total, our analysis provides a comprehensive resource for biomarker and therapeutic target identification, and suggests that copy number and methylation profiling should complement tumor sequencing efforts to improve patient risk assessment.

## Introduction

Cancers that arise from the same tissue can exhibit vast differences in clinical behavior. For instance, among individuals diagnosed with early-stage colorectal cancer, about 60% of patients will be cured by surgery alone, while the remaining 40% will experience a recurrence that is frequently fatal (*1*). Standard methods to predict patient risk rely on clinical and pathological evaluation of the primary tumor. However, blinded studies have revealed low levels of concordance among pathologists assessing the same tumor (*2*–*5*), and even perfect cancer staging provides limited prognostic information on a patient’s likely outcome (*6*–*9*).Improved methods to stratify patient risk could aid clinical decision-making and identify the cancer patients most likely to benefit from surveillance, surgery, or chemotherapy.

Advances in high-throughput technologies have yielded unprecedented insight into the diverse array of genomic changes found within every cancer cell. Projects like The Cancer Genome Atlas (TCGA) (*10*) and the International Cancer Genome Consortium (ICGC) (*11*) have characterized methylation, mutation, copy number, and gene expression patterns across cancer types. As a result of these studies, many of the genomic differences between normal and transformed cells have been identified and characterized. However, we lack a similar understanding of the genomic differences between benign tumors, which can be left untreated or cured with surgery alone, and aggressive tumors, which recur after treatment and often prove deadly. Furthermore, while metastatic dissemination is the ultimate cause of 90% of cancer deaths (*12*), no metastasis-specific therapies have been successfully applied in the clinic. The discovery of genomic alterations found only in aggressive tumors could prompt the development of new therapies that can be applied to specifically block the cancer phenotypes usually responsible for patient death.

Previous efforts to discover molecular markers of deadly cancers have largely focused on the gene expression changes associated with patient prognosis (*13*–*16*). These studies have identified a set of transcripts that encode proteins involved in cell cycle progression that correlate with recurrence and death in several cancer types (*13*, *17*–*22*). However, it is unknown whether these genes convey prognostic information not already captured by common markers of tumor proliferation, like Ki-67 staining (*23*). Additionally, targeted and whole-genome tumor sequencing is becoming increasingly routine at many major hospitals (*24*, *25*). These sequencing efforts can identify molecular alterations, like *BRAF* or *EGFR* mutations, that confer sensitivity to particular therapies, and could shed light on tumor progression (*26*). However, no large-scale studies have tested the prognostic value of genome-wide sequencing, or any of the other types of genomic analyses that are now possible. It is not clear whether genomic alterations can accurately stratify patient risk, and, if they can, whether different cancer types exhibit unique or common genomic signatures of fatal disease. Finally, the majority of cancer genomic profiles that have been collected to date are spread across different databases and formats, further complicating efforts to comprehensively analyze patient outcome.

In order to gain a global understanding of the genomic features in a primary tumor that influence cancer prognosis, we collected and analyzed molecular profiles from 33,036 patients with solid tumors. We report that single base-pair mutations in cancer driver genes are almost always unlinked with patient survival, while copy number, methylation, and gene expression signatures provide significant prognostic information across cancer types. Our comprehensive, gene-centric analysis sheds light on the genomic changes that drive aggressive disease and will provide a useful resource for the development of prognostic biomarkers to improve patient stratification.

## A cross-platform, pan-cancer analysis of cancer survival data

To determine the differences between benign and fatal tumors, we first analyzed multiple classes of genomic data from 9,442 patients with 16 types of cancer from the TCGA (outlined in Figure S1A; abbreviations are defined in Figure S1B). For every tumor type and every dataset, we generated Cox univariate proportional hazards models linking the presence or expression of a particular feature with clinical outcome. We report the Z score for each model, which encodes both the directionality and significance of a particular association. If the presence of a mutation or the expression of a gene is significantly associated with patient death, then a Z score >1.96 corresponds to a P value <.05 (Figure S2A-C). In contrast, a Z score less than −1.96 indicates that the presence of a mutation or the expression of a gene is significantly associated with patient survival. To identify genomic features with pan-cancer prognostic significance, Z scores from individual studies can also be combined using Stouffer’s method (see Materials and Methods).

We extracted mutation, copy number, methylation, gene expression, and clinical information from 16 TCGA cohorts (summarized in Table S1). To assess the validity of our data analysis pipeline, as well as the accuracy of the reported patient annotations, we first examined the overall survival curves for the 16 tumor types that we profiled. As expected, we observed significant differences in clinical outcome according to a cancer’s tissue-of-origin (Figure S2C).Prostate cancer had the least aggressive clinical course, with a median survival time that was not reached in this dataset (>4600 days), while pancreatic cancer conferred the most dismal prognosis (median survival time: 444 days). Overall, the 5-year survival frequencies of patients in the TCGA were highly similar to the national averages reported by NCI-SEER (*27*), suggesting that the patients included in this analysis are broadly representative of the general population (Figure S2D). Next, we inferred patient sex on the basis of chromosome-specific gene expression patterns (*13*, *28*). Our analysis exhibited >99% concordance with patients’ self-reported sex, further verifying the overall accuracy of the clinical annotations and our data processing pipeline (Figure S2E).

## Cancer mutations convey limited prognostic information

We first set out to discover whether single base-pair mutations in cancer genomes were associated with patient outcome. We identified non-silent mutations in each tumor, and then compared survival times for patients harboring wild-type copies of a gene to patients that harbored a mutation in that gene. Surprisingly, our analysis uncovered very few mutations that were significantly associated with patient survival (Figure 1 and Table S2A). We first focused on known oncogenes and tumor suppressors, and found that among the 30 most-frequently mutated cancer driver genes, only two (*EGFR* and *TP53*) were associated with outcome in more than two tumor types (Figure 1A). *TP53* mutations were linked to outcome in five of 16 cancer types, though the differences in patient survival were generally small (e.g., a median survival of 3736 days vs. 3430 days for *TP53*-WT and *TP53*-mutant breast cancer patients, respectively; Figure S3A). In contrast, many other cancer driver genes were not associated with survival time in any tumor type. While mutations in *KRAS*, *PIK3CA*, *CDKN2A*, *BRAF*, *KMT2D*, *ATM*, *SMAD4*, and many other genes were frequently observed, they were never significantly linked with patient outcome.

We next tested several alternate approaches to discover prognostic mutations. We identified recurrently-mutated codons in tumor genomes, and asked whether mutations in these specific residues conveyed prognostic information. IDH1-c132 mutations were significantly associated with a favorable prognosis in glioma, but other “hotspot” mutations (KRAS-c12, PIK3CA-c1047, BRAF-c600, etc.) were largely uninformative (Figure S4A). Pooling all “hotspot” mutations per gene together also failed to uncover robust survival associations (Figure S4B-C).We then asked whether the presence of mutations in multiple cancer driver genes might cooperate to confer a worse clinical outcome. We found that, in general, patients harboring mutations in two cancer driver genes that were not prognostic alone had the same risk of death as patients with wild-type copies of one or both genes (Figure S4E). Next, we considered the possibility that the clonality of a mutation might affect its prognostic significance. We calculated the variant allele frequency (VAF) for each cancer mutation, and tested whether mutations present at higher levels in single tumors were more likely to be associated with outcome. We found that restricting our analysis to mutations with high VAFs failed to identify more prognostic genes, indicating that patient stratification is unlikely to be improved by assessing only clonal mutations (Figure S5).

**Figure 1.**
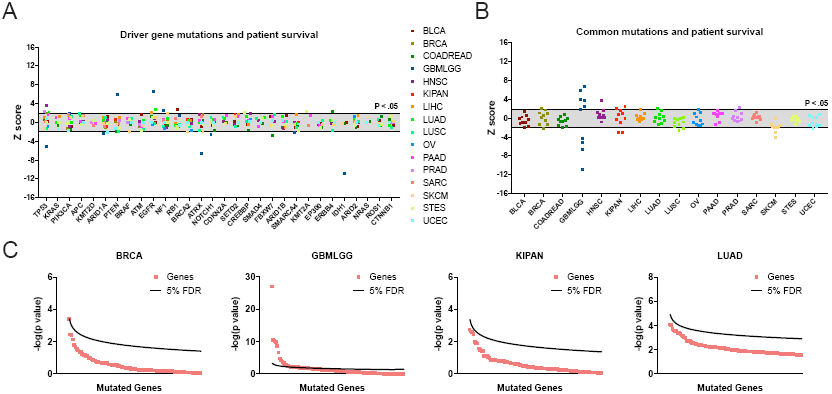
Single base-pair mutations convey limited prognostic information. (A) Z scores of the 30 most frequently-mutated (*29*) cancer driver genes in 16 tumor types from the TCGA are displayed. Z scores were calculated by regressing survival times between patients harboring wild-type and mutant copies of a gene if a gene was mutated in ≥2% of samples per tumor type. Dotted lines are plotted at Z = ±1.96, corresponding to P < .05. The complete list of Z scores are presented in Table S2A. (B) Z scores of the 10 most frequently-mutated genes per cancer type are displayed. Dotted lines are plotted at Z = ±1.96, corresponding to P < .05. (C) Univariate Cox models for all genes mutated in ≥2% of patients in a particular tumor type were determined and then sorted according to P value. The P values of the top-scoring genes are plotted against a line corresponding to a 5% false-discovery rate. While several genes are significant at this threshold in glioma, very few significant genes were found in other cancer types.

Finally, we investigated whether mutations in genes other than known oncogenes and tumor suppressors might affect prognosis. We therefore expanded our analysis to include all genes mutated in ≥2% of patients with a particular tumor type. To account for greatly expanding the number of genes tested, we applied a Benjamini-Hochberg correction with a 5% false-discovery rate to the individual Z scores that we obtained. We uncovered several genes that were linked with prognosis in glioma, but found very few genes significantly associated with death or survival in the other 15 cancer types (Figure 1B-C and Table S2A). For instance, in breast cancer and lung adenocarcinoma, 128 and 3,996 genes were mutated in ≥2% of patients, respectively, but none of these mutations were significantly correlated with patient outcome at a 5% FDR. These results suggest that non-oncogenic mutations tend to be no more prognostic than mutations in cancer driver genes.

In our above analysis, we noted that the five genes with the strongest survival associations were observed in the GBMLGG cohort. As glioma appeared to be an exception to our overall finding that mutations are rarely prognostic, we decided to investigate this cancer type further. Among the top-scoring genes, we found that *PTEN* and *EGFR* mutations conferred dismal prognosis, while mutations in *IDH1*, *TP53*, and *ATRX* were associated with favorable prognosis (Figure S6A). Mutations in these genes have previously been linked to distinct glioma subtypes (*30*–*32*), and we verified that mutations in *IDH1*, *TP53*, and *ATRX* were most frequently observed in low-grade gliomas, while mutations in *PTEN* and *EGFR* were most frequently observed in glioblastomas (Figure S6B). The glioma subtypes exhibited distinct clinical courses that were largely able to explain the differences in prognosis conferred by these mutations (Figure S6C-D). Interestingly, we observed a set of patients with subtype-discordant mutations (e.g., *PTEN* or *EGFR* mutations in low-grade glioma or *IDH1* mutations in glioblastoma). Analysis of GBM-distinctive chromosome copy number changes (*33*) suggested that these mutation-discordant tumors had been misclassified pathologically, and that the *PTEN* and *EGFR*-mutant LGGs were in fact glioblastomas, and vice-versa (Figure S6E). Consistent with this hypothesis, we found that patients with low-grade glioma who had mutations in *PTEN* or *EGFR* had a significantly worse outcome compared to other LGG patients (median survival of 315 days vs. 2379 days, Figure S6F). Thus, in the case of glioma, DNA mutation analysis captures subtype-specific survival patterns and can potentially correct pathological misdiagnosis. However, outside of glioma and the tumor suppressor *TP53*, single base-pair mutations rarely convey prognostic information.

## Oncogene and tumor suppressor CNAs are associated with cancer patient mortality

As single base-pair mutations were largely uninformative, we next set out to determine whether gene copy number conveyed prognostic information. We regressed gene copy number, as measured by SNP arrays, against patient outcome in each individual tumor type. We then examined the clinical impact of CNAs affecting the same 30 cancer driver genes that we previously investigated. Surprisingly, we found that the copy number of these oncogenes and tumor suppressors was frequently linked with patient outcome (Figure 2 and Table S2B).Amplification of *EGFR*, *PIK3CA*, and *BRAF*, and deletion of *CDKN2A*, *RB1* and *EP300* were strongly associated with shorter patient survival times in four or more cancer types each. Copy number was prognostic even for genes in which mutations were not linked with outcome: for instance, while mutations in *PIK3CA* were never informative, the copy number of *PIK3CA* was associated with outcome in breast, colorectal, glioma, lung-squamous, pancreas, and prostate cancers. Overall, among the 30 most frequently-mutated cancer driver genes, we detected 108 significant associations between gene copy number and outcome, compared to 23 associations between mutation and outcome. For 28 out of 30 driver genes, DNA copy number was prognostic in more cancer types than mutational status was (Table 1). We conclude that determining the copy number of oncogenes and tumor suppressors in a primary tumor can better stratify patient risk than assessing single base-pair mutations alone.

**Figure 2.**
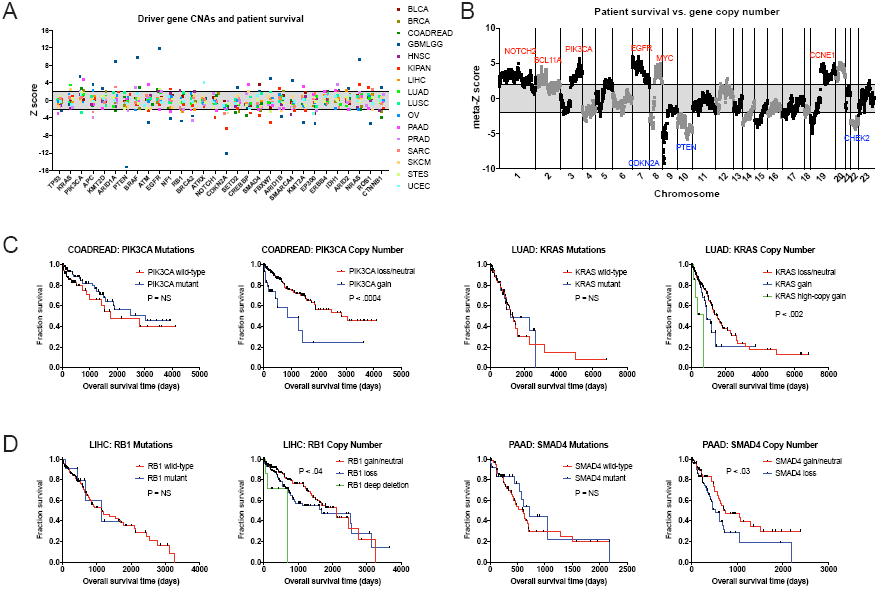
Oncogene and tumor suppressor CNAs drive cancer patient mortality. (A) Z scores regressing gene copy number against patient outcome for the 30 most frequently-mutated (*29*) cancer driver genes are displayed. Dotted lines are plotted at Z = ±1.96, corresponding to P < .05. The complete list of Z scores are presented in Table S2B. (B) Z scores from 16 cancer types from the TCGA were combined using Stouffer’s method, and then the resulting meta-Z scores were plotted against the chromosomal location. Genes were binned by average Z score into groups of 5 for visualization. (C) Kaplan-Meier curves are plotted for two oncogenes, *PIK3CA* (left) and *KRAS* (right), comparing the prognostic relevance of mutations in these genes versus copy number alterations in these genes. (D) Kaplan-Meier curves are plotted for two tumor suppressors, *RB1* (left) and *SMAD4* (right), comparing the prognostic relevance of mutations in these genes versus copy number alterations in these genes.

**Table 1.**

| Gene | Mutations |  | CNAs |  |
| --- | --- | --- | --- | --- |
|  | # Cancer<br>Types P < .05 | Max Abs(Z) | # Cancer<br>Types P < .05 | Max Abs(Z) |
| TP53 | 5 | -5.15 | 2 | -2.92 |
| KRAS | 0 | -1.56 | 2 | 3.53 |
| PIK3CA | 0 | 1.91 | 6 | 5.53 |
| APC | 1 | 1.97 | 4 | -3.92 |
| KMT2D | 0 | -0.62 | 2 | 2.70 |
| ARID1A | 2 | -2.31 | 3 | 8.82 |
| PTEN | 1 | 5.96 | 2 | -15.16 |
| BRAF | 0 | -1.89 | 4 | 9.76 |
| ATM | 0 | -1.77 | 3 | -5.09 |
| EGFR | 3 | 6.66 | 6 | 11.86 |
| NF1 | 1 | 2.46 | 3 | -4.19 |
| RB1 | 1 | 2.77 | 5 | -4.70 |
| BRCA2 | 1 | -2.03 | 5 | -4.48 |
| ATRX | 1 | -6.57 | 1 | 4.03 |
| NOTCH1 | 1 | -2.51 | 2 | -3.80 |
| CDKN2A | 0 | 1.50 | 6 | -12.10 |
| SETD2 | 0 | 1.75 | 3 | 3.00 |
| CREBBP | 1 | 2.10 | 4 | 3.66 |
| SMAD4 | 0 | 0.98 | 5 | 3.68 |
| FBXW7 | 1 | -2.82 | 3 | 5.03 |
| ARID1B | 0 | 1.45 | 4 | -3.71 |
| SMARCA4 | 1 | -2.21 | 3 | 4.58 |
| KMT2A | 1 | 2.37 | 3 | -4.95 |
| EP300 | 0 | -1.06 | 4 | -5.20 |
| ERBB4 | 1 | 2.42 | 3 | 2.15 |
| IDH1 | 1 | -10.93 | 4 | 3.65 |
| ARID2 | 0 | 1.47 | 4 | 2.68 |
| NRAS | 0 | 1.16 | 4 | 9.40 |
| ROS1 | 0 | 1.26 | 4 | -5.17 |
| CTNNB1 | 0 | 1.21 | 4 | -3.82 |

We next investigated whether these oncogenic CNAs were likely to be drivers of this phenomenon, or whether they were passenger genes that changed in copy number along with other, unknown drivers. To assess this question, we combined Z scores from different cancer types using Stouffer’s method, and then plotted the pan-cancer meta-Z scores along every chromosome (Figure 2B). This analysis revealed multiple sharp peaks and valleys in the data that overlapped with known driver mutations. The most significant survival-associated copy number changes genome-wide were found on chromosome 9p in a valley that precisely included the tumor suppressor *CDKN2A*. Z score peaks were found at loci that include oncogenes *PIK3CA*, *EGFR*, *MYC*, *CCNE1*, and others. This overlap suggests that, in many instances, the copy number of these oncogenes and tumor suppressors are directly influencing the risk of cancer patient death.

We considered one alternate explanation of this data. The tumor microenvironment contains a mixture of cell types, including endothelia, stroma, and immune cells. We reasoned that the presence of euploid immune cells could blunt our ability to detect copy number alterations, and a robust immune infiltrate could itself signify a favorable prognosis. Thus, in this model, copy number alterations are not causally linked with patient outcome, but instead indicate poor prognosis due to the lack of an immune response. However, three lines of evidence argue against this interpretation. First, in 15 of 16 cancer types, tumor purity is not associated with patient outcome (Figure S7A-B). Secondly, in multivariate Cox models that include gene copy number and tumor purity, the copy number of cancer driver genes remain broadly prognostic, indicating that CNAs are associated with outcome even in highly-pure tumor samples that lack significant immune infiltration (Figure S7C-D). Finally, if CNAs were prognostic due to the lack of a euploid cell infiltrate, then this would imply that the presence of most copy number alterations would be prognostic. However, in multiple cancer types, we found that recurrent CNAs and survival-associated CNAs encompass different genes (Figure S7E). For instance, in colorectal cancer, the 20q13 locus is amplified in ∽51% of patients, but is not associated with dismal prognosis (Figure S7F). In total, these data suggest that cancer copy number alterations are independent drivers of patient mortality, and do not simply reflect the absence of an immune response.

## Methylation of developmental transcription factors is associated with poor prognosis across cancer types

Cancers exhibit global alterations in the level of CpG methylation, a nucleotide modification that typically suppresses transcription (*34*). However, the prognostic importance of tumor methylation patterns across cancer types is unknown. To test whether methylation could stratify cancer patient risk, we regressed the mean β-value (methylation level) per gene in every tumor in the TCGA cohorts against patient outcome. We discovered a median of 2,237 genes per tumor type significantly associated with outcome (Figure 3A-D and Table S2C). To find genes with pan-cancer prognostic power, we applied Stouffer’s method to combine Z scores across cancer types. Applying a 5% FDR correction and generating meta-Z scores from ∽1,000,000 random permutations of the data verified that hundreds of genes were more strongly associated with patient outcome across cancer types than expected by chance (Figure 3B-C). Gene ontology analysis of methylated loci in tumors with grim prognosis revealed a striking enrichment of transcription factors and genes involved in embryonic development, including *NKX6-1*, *HOXD12*, and *FOXE1* (Figure 3D and Table S3A-B). In contrast, fewer functional categories were significantly enriched among genes associated with favorable prognosis, though the most enriched GO category in this gene set was olfactory receptor activity.

**Figure 3.**
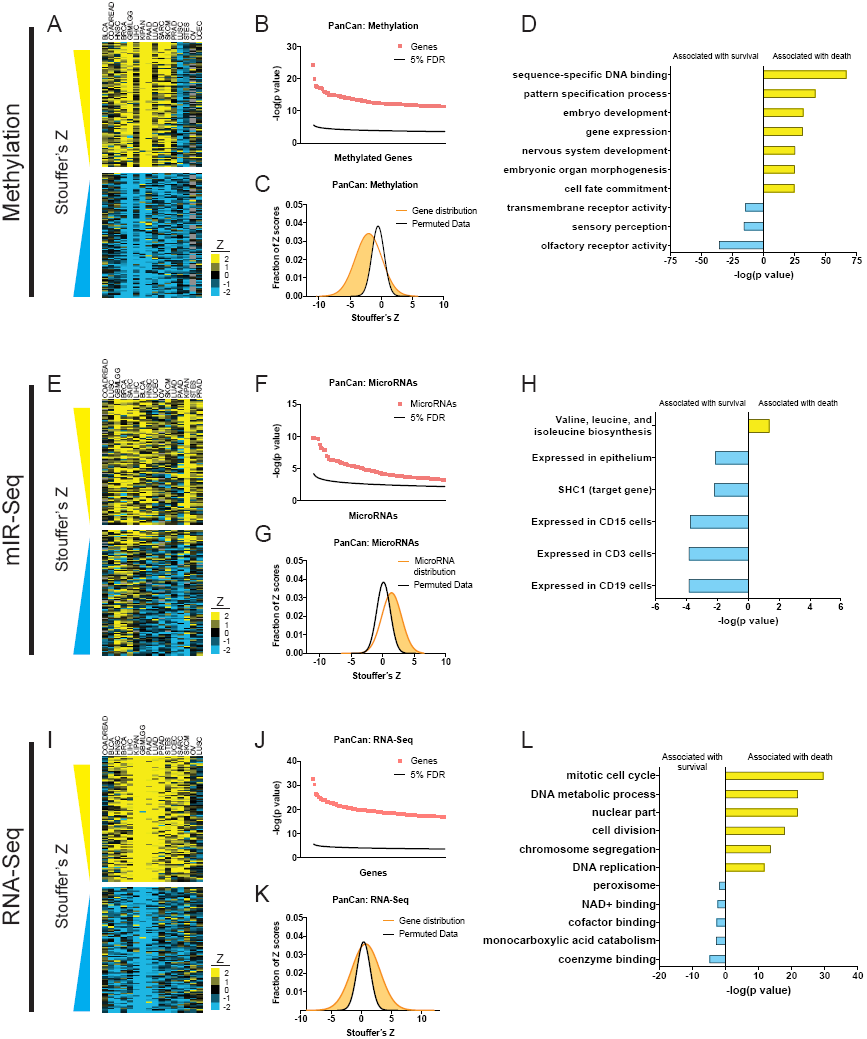
Methylation, microRNA, and gene expression patterns in primary tumors are correlated with patient outcome. (A, E, and I) Z scores from individual cancer types were combined using Stouffer’s method, and a heatmap of the 100 methylation sites (A), microRNAs (E) or genes (I) with the largest or smallest meta-Z scores are displayed. The complete list of Z scores are presented in Table S2C-E. (B, F, and J) The significance of the top-scoring methylation sites (B), microRNAs (F), or genes (J) are plotted against a 5% false-discovery threshold calculated using the Benjamini-Hochberg method. 9,774 methylation sites, 218 microRNAs, and 7,023 genes exhibit a significant correlation with patient outcome across cancer types at a 5% false-discovery threshold. (C, G, and K) A histogram of meta-Z scores for methylation sites (C), microRNAs (G), or genes (K) are plotted against a histogram of meta-Z scores from ∽1,000,000 random permutations of the underlying patient data. (D, H, and L) Gene ontology terms over-represented among methylation sites (D), microRNAs (H), or genes (L) that are significantly associated with either patient survival or patient death are displayed. The complete lists of GO terms are presented in Table S3.

Previous research has demonstrated that cancers exhibit *de novo* methylation of developmental genes that are silenced by the chromatin-modifying Polycomb complex during embryogenesis (*35*). These Polycomb targets include lineage-defining transcription factors that are activated or repressed to specify tissue identity (*36*). Our finding that developmental transcription factors like *NKX6-1* and *HOXD12* were methylated in aggressive tumors led us to inquire whether death-associated methylation sites were also linked with Polycomb activity.Indeed, we observed a highly-significant enrichment of Polycomb component Suz12 binding sites among the genes where increasing methylation conferred shorter patient survival (Figure S8A). We further found that methylation of the Suz12-associated loci blocked transcription of these genes, and, across tumors, high levels of Suz12-associated methylation was frequently observed in high-grade (de-differentiated) malignancies (Figure S8B-C). This data suggests that cancers methylate and silence lineage-defining transcription factors, which promotes loss of differentiation and is associated with aggressive disease. In contrast, the olfactory genes associated with patient survival were barely expressed in any tumor type, and this low level of transcription was unaffected by methylation (Figure S8D-F). Whether the methylation of olfactory receptors plays any direct role in patient outcome is at present unknown.

## A microRNA-SHC1 circuit influencing patient survival

Using our survival analysis pipeline, we next analyzed mir-Seq data from the TCGA cohorts. Out of 830 microRNAs detected in 12 or more tumor types, 218 were significantly associated with patient death or survival across cancer types (Figure 3E-H and Table S3C-D).Gene set enrichment analysis revealed a slight overrepresentation of microRNAs involved in branched-chain amino acid metabolism among microRNAs associated with poor prognosis.MicroRNAs associated with survival were enriched for those expressed in CD3+, CD15+, or CD19+ cells as well as a set of microRNAs that target the receptor-tyrosine kinase adaptor protein *SHC1* (*37*) (Figure 3H). Across tumor types, we found that high expression of the *SHC1*-targeting microRNAs was associated with lower levels of SHC1 expression (Figure S9A).Increasing *SHC1* expression and decreasing *SHC1*-targeting microRNA expression were associated with higher levels of MEK1 phosphorylation, a downstream readout of RTK signaling (Figure S9B-C). We also observed a strong anti-correlation between the prognostic power of *SHC1* expression and the *SHC1*-targeting microRNAs: in cancer types where high *SHC1* expression was deadly, high expression of the *SHC1*-targeting microRNAs was protective, and vice-versa (Figure S9D). Finally, in multivariate proportional hazards models, accounting for *SHC1* expression decreased the prognostic significance of the *SHC1*-targeting microRNAs (Figure S9E-H). In total, our analysis demonstrates that a large fraction of expressed microRNAs capture prognostic information across cancer types, and a set of microRNAs associated with patient survival may exert this function by dampening the output of receptor tyrosine kinase signaling.

## High expression of cell cycle genes at the RNA or protein level indicates grim prognosis

As a final set of primary data to examine, we considered gene expression in tumors as measured by RNA-Seq and reverse-phase protein arrays (RPPAs). Regressing RNA-seq values against patient outcome identified 7,023 genes significantly associated with clinical prognosis within and across cancer types (Figure 3I-L and Table S2E). Previous pan-cancer analyses have reported that the expression of genes related to cell cycle progression indicates poor prognosis (*13*), and our GO analysis of death-associated factors revealed a strong enrichment of genes involved in mitosis and DNA metabolism (Figure 3L and Table S3E-F).Indeed, the expression levels of top-scoring genes were strongly correlated with the expression of proliferation marker *MKI67* in normal tissue, suggesting that most of these genes capture the rate of cell division within a tumor (Figure S10A-B). We observed a similar pattern among the smaller set of proteins and protein modifications that had been measured using RPPAs (Figure S10C-E). The expression of cyclin B1, EGFR, and the phosphorylated form of Rb1 were among the top-scoring genes associated with death from multiple cancer types (Table S2F). Overall, there was a strong correlation between the Z scores calculated from RNA-Seq and RPPAs (R = 0.54; Figure S10E). This indicates that survival-associated gene expression patterns discovered via RNA profiling are present at the protein level as well, and suggests that many of the prognostic genes we have identified can be assessed via immunohistochemistry for easier clinical adoption.

## Independent patient cohorts verify the prognostic significance of cancer-associated CNAs and cell cycle transcripts

To confirm the generality of our findings, we collected independent patient cohorts harboring mutation, copy number, or gene expression data linked to survival outcome. We first examined a set of 16 patient cohorts from the International Cancer Genome Consortium (ICGC) comprising 3,310 patients with whole-genome or whole-exome sequencing, and utilized our survival analysis pipeline to identify outcome-associated mutations. Overall, the mutation frequencies and the Z scores of common mutations were highly similar between the ICGC and TCGA cohorts (R = 0.67, P < .0001, and R = 0.35, P < .002, respectively; Figure 4A-B).Consistent with our TCGA analysis, mutations in *TP53* were associated with outcome in more patient cohorts than any other mutation, though the differences in median survival times were small (Figure 4C and S3B). Other mutations, including in known cancer driver genes, were rarely associated with outcome in individual cancer types and harbored minimal pan-cancer significance (Figure 4D-E and Table S4A). Mutations in *KRAS*, *PIK3CA*, *BRAF*, *APC*, *PTEN*, *CDKN2A*, and many others were frequently observed, but were never correlated with outcome (Figure 4D). In contrast, among 2,398 patients with CNA data collected from cBioportal, survival analysis revealed numerous amplifications and deletions associated with patient mortality across cancer types (Table S4B). We found that prognostic amplifications were centered around oncogenes, including *TERT*, *MYC*, and *MDM2*, while prognostic deletions encompassed tumor suppressors *CDKN2A*, *PTEN*, and *TP53* (Figure 4F). Overall, we observed a highly significant correlation between the meta-Z scores obtained from cBioportal and the corresponding TCGA cohorts (R = 0.54; Figure 4G). Finally, we collected a set of 17,886 patients from the Gene Expression Omnibus (GEO) with microarray data and linked clinical outcomes. Death-associated genes were strongly enriched for biological processes associated with mitotic progression, and Z scores for individual genes were highly related between the independent TCGA and GEO cohorts (R = 0.67; Figure 4H-I). In total, these analyses suggest that the survival patterns discovered in the TCGA dataset are in fact common across cancer patients. In particular, while mutations in genes other than *TP53* are typically non-prognostic, copy number alterations in known cancer driver genes and the expression levels of mitotic cell cycle factors strongly influence patient mortality.

**Figure 4.**
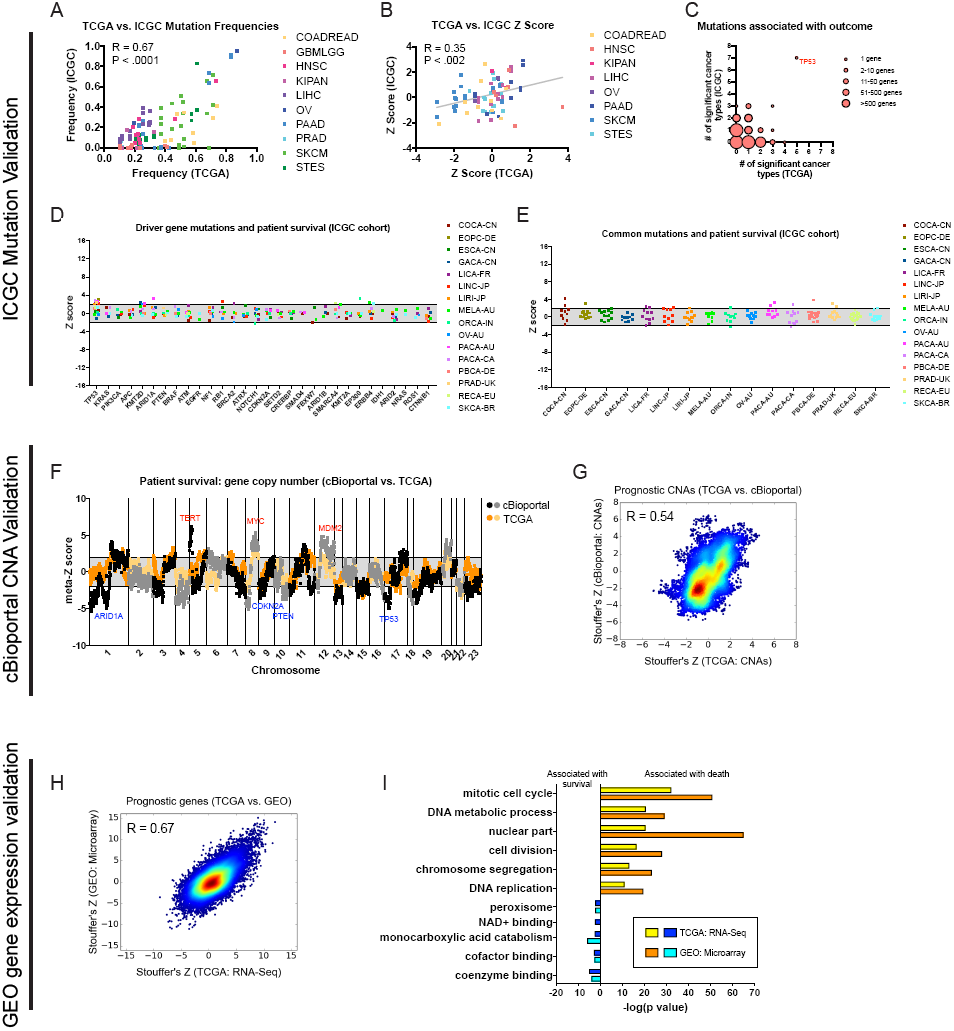
Driver gene copy number and cell cycle transcripts, but not driver gene mutational load, are associated with survival in independent patient cohorts. (A) Genes mutated in ≥10% of patients in each tumor type in the TCGA were identified, and then compared to the mutation frequency of these genes in the corresponding ICGC cohort or cohorts. The complete list of Z scores are presented in Table S4A. (B) Z scores of the 10 most frequently-mutated genes per cancer type in the ICGC were identified and then plotted against the Z scores of the same gene from the corresponding TCGA cohort or cohorts. (C) Significant Z scores (>1.96 or <-1.96) were counted per gene, and then the number of significant cohorts from the TCGA and the ICGC are plotted. While the vast majority of frequently-mutated genes are significant in zero or one cancer type, *TP53* mutation status is associated with prognosis in 12 of 32 patient cohorts. (D) Z scores of 30 cancer driver genes in 16 tumor types from the ICGC are displayed. Z scores were calculated comparing survival times between patients harboring wild-type and mutant copies of a gene for all genes mutated in ≥2% of samples per tumor type. Dotted lines are plotted at Z = ±1.96, corresponding to P < .05. (E) Z scores of the 10 most frequently-mutated genes per cancer type in the ICGC are displayed. Dotted lines are plotted at Z = ±1.96, corresponding to P < .05. (F) Z scores for the copy number of each gene from four cancer types from cBioportal or from the corresponding four TCGA cancer types (BLCA, BRCA, LUAD, and PRAD) were combined using Stouffer’s method. The resulting meta-Z scores were plotted against the chromosomal location, binned every five genes for visualization. The full list of Z scores are presented in Table S4B. (G) Meta-Z scores from TCGA are plotted directly against meta-Z scores from the corresponding four cancer types from cBioportal. (H) Meta-Z scores from TCGA (RNA-Seq) are plotted against meta-Z scores from the GEO (microarray) patient cohorts. The complete list of Z scores by cancer type in GEO are presented in Table S4C. (I) Gene Ontology terms over-represented among death‐ or survival-associated transcripts from the TCGA and GEO analysis are plotted. Complete GO term lists are presented in Table S3E-F and S4D-E.

## Cross-platform analysis of cancer survival factors

Given the ability to assess any genomic feature in a primary tumor, what measurement confers the most prognostic information? We found that, across platforms and cancer types, the expression of the mitotic kinesin *KIF20A* and the centrosomal kinase *PLK1* exhibited the strongest association with cancer death, while methylation of the gap junction component *PANX1* and copy number of the tumor suppressor *CDKN2A* displayed the most significant associations with survival (Figure 5A-B). Strikingly, 99 out of the top 100 death-associated features were mRNA transcripts, while top-scoring survival-associated features included a mix of CNA’s, transcripts, and methylation sites. These patterns were largely recapitulated among the top-scoring genes within single cancer types (Figure S11A-B). No cancer mutations were among the top 500 features associated with death or survival across cancers, and mutations made up less than 2% of top-scoring features within single cancer types, further demonstrating that other categories of genomic data are superior indicators of patient prognosis.

**Figure 5.**
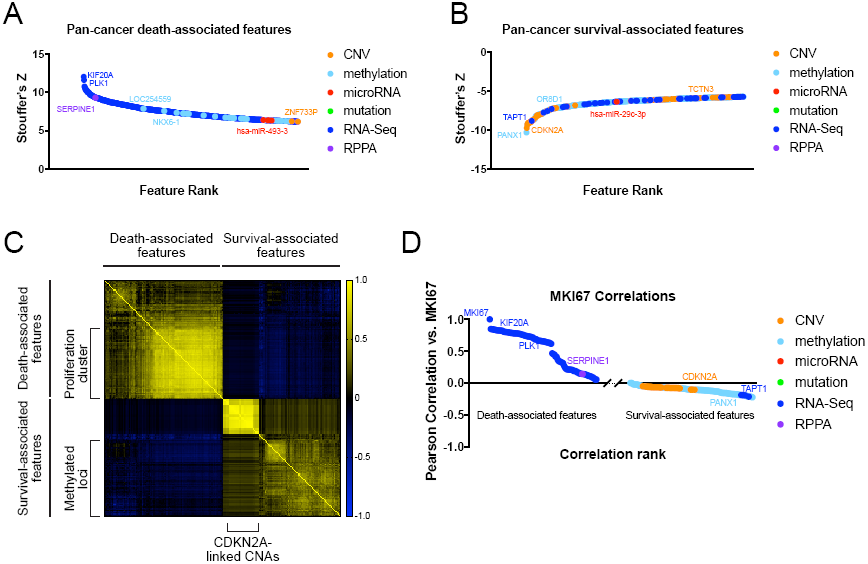
Cross-platform analysis of prognostic features. (A) Meta-Z scores of the top 1% of genomic features associated with death from cancer are displayed, color-coded according to feature type. (B) Meta-Z scores of the top 1% of genomic features associated with cancer survival are displayed, color-coded according to feature type. (C) A heatmap of pairwise Pearson correlations of the top 100 features associated with either death or survival are displayed. Clusters of methylation sites, *CDKN2A*-linked CNAs, and proliferation genes are indicated. Blue and yellow colors indicate the strength of the pairwise correlation. (D) The top 100 features associated with either death or survival are displayed, sorted by their correlation score with the proliferation marker *MKI67*.

Individual features can potentially be combined to build multivariate models with improved stratification power. However, prognostic variables that are interrelated fail to provide any additive benefit. We therefore sought to discover whether the top-scoring outcome-associated features captured independent aspects of cancer biology, or whether they represented shared measurements of underlying phenotypes. Indeed, a correlation matrix of the top-scoring features revealed strong similarities across individual tumors in the expression values among most death-associated transcripts (Figure 5C). The largest cluster consisted of genes with well-described roles in cell-cycle progression, including *CDC20*, *CCNB1*, and *AURKA*, and exhibited strong individual correlations with the proliferation marker *MKI67* (Figure 5D). Survival-associated CNAs were also highly interrelated, reflecting their shared chromosomal linkage with *CDKN2A*, while survival-associated methylation sites exhibited less significant correlations. Interestingly, the survival-associated features were only mildly anti-correlated with *MKI67*, suggesting that they did not primarily identify tumors with low levels of mitotic activity. In total, this analysis suggests that many death-associated transcripts are potentially redundant reporters of tumor proliferation.

## Multivariate models with a cell cycle meta-gene improve patient risk prediction and provide insight into tumor biology

To identify genomic features that correlate with patient survival and are independent of tumor mitotic activity, we constructed multivariate Cox models using a previously-described cell cycle meta-gene, PCNA25 (*38*) (Figure 10A and Table S5). This meta-gene consists of the 25 genes that exhibit the strongest co-expression pattern with the proliferation marker *PCNA*, and we found that PCNA25 expression correlates with the doubling time of cancer cell lines *in vitro* and with Ki-67 staining *in vivo* (Figure S12B-C). Controlling for cell proliferation with PCNA25 resulted in a remarkably different set of transcripts associated with death from cancer (Figure 6A-B). Though *KIF20A* was not a component of the PCNA25 meta-gene, in multivariate models, *KIF20A* dropped from the top-ranked gene to the 645^th^–ranked gene (Table S5E). The top-ranked genes with PCNA25 were the protease inhibitor *SERPINE1* and the extracellular crosslinking enzyme *LOX*, and GO term analysis revealed a striking enrichment of extracellular, cell motility, and angiogenesis factors among death-associated genes (Figure 6B and Table S6). PCNA25 further decreased the prognostic power of most cancer mutations; for instance, after controlling for proliferation, *TP53* mutations were significant in only two cancer types (HNSC and GBMLGG; Figure S12D and Table S5A). However, many CNAs and methylated genes remained prognostic, particularly deletions that included *CDKN2A* and the methylation of developmental transcription factors (Figure 10E-F and Table S6). In individual tumor types, the top death-associated features after PCNA25 correction tended to be mRNA transcripts, while survival-associated features included CNAs, transcripts, and methylation sites (Figure S11C-D).

**Figure 6.**
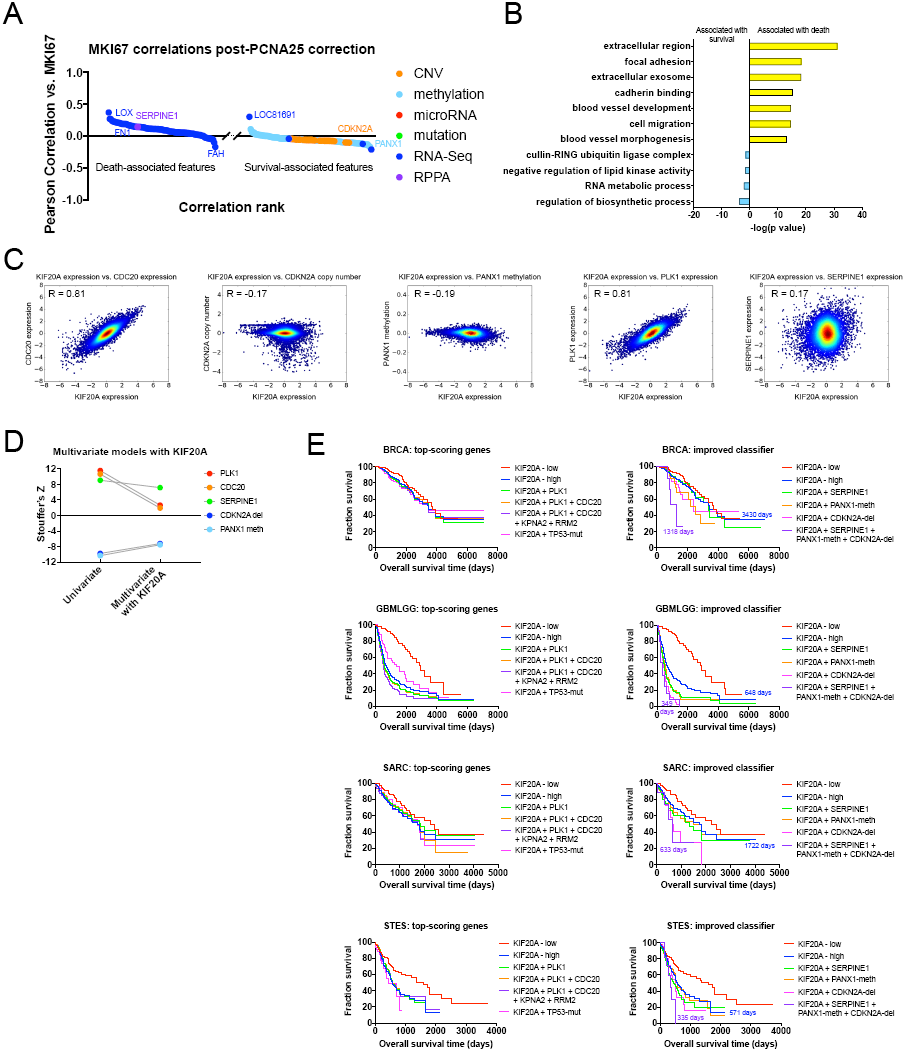
Multivariate analysis with the PCNA25 meta-gene identifies proliferation-independent prognostic features. (A) The top 100 features in multivariate Cox proportional hazards models with PCNA25, associated with either death or survival, are displayed. The features are sorted by their correlation coefficient with the proliferation marker *MKI67*. (B) Gene ontology terms over-represented among transcripts that are significantly associated with either patient survival or patient death, after PCNA25 correction, are displayed. The complete lists of GO terms are presented in Table S6. (C) Density plots displaying the pairwise correlation between *KIF20A* expression and *PLK1* expression, *CDC20* expression, *SERPINE1* expression, *CDKN2A* copy number, or *PANX1* methylation. (D) Multivariate models were constructed containing *KIF20A* expression and each genomic feature presented in Figure 6C. Pan-cancer Stouffer’s Z scores were calculated across the 16 TCGA cohorts, and univariate vs. multivariate Z scores are plotted. (E) Kaplan-Meier curves of various combinations of genomic features are displayed. The left plots display survival curves generated by the sequential addition of top-scoring proliferative features (*KIF20A, PLK1, CDC20, KPN2A, RRM2*), split based on the average expression of the indicated gene or genes. For ease of visualization, only tumors with below-average *KIF20A* are displayed, every other survival curve is for tumors with above-average expression. The right plots display survival curves generated by the sequential addition of features that remained prognostic after PCNA25 correction. PANX1-meth indicates patient with below-average levels of PANX1 methylation, while CDKN2A-del indicates patients with CDKN2A deletions (copy number < −0.3). Numbers on each graph indicate the median survival times of the KIF20A-high tumors (blue) and tumors identified with the combined classifier (purple).

To test whether controlling for proliferation could improve patient stratification, we constructed multi-gene models harboring various top-scoring genomic features. Across all tumors, *KIF20A* was highly correlated with *PLK1* and *CDC20*, the second‐ and third-highest scoring genes in the proliferation cluster, while *KIF20A* was marginally correlated or anti-correlated with *CDKN2A* copy number, *PANX1* methylation, and *SERPINE1* expression, three features identified after PCNA25 correction (Figure 6C). Accordingly, in multivariate models containing *KIF20A*, *PLK1* and *CDC20* expression lost their prognostic significance in most cancer types, while models containing *CDKN2A*, *PANX1*, or *SERPINE1* retained significance in five to eight cancer types each (Figure 6D). To further explore this observation, we generated Kaplan-Meier curves using various combinations of proliferation-dependent and ‐independent features. We found that a single proliferation marker (*KIF20A*) identified at-risk patients with nearly the same efficacy as models that combined several of the top-scoring proliferation-associated genes (Figure 6E). Similarly, assessing *TP53* mutation status generally failed to improve on the stratification conferred by *KIF20A* expression alone. In contrast, when we combined *KIF20A* expression with various proliferation-independent features, including *SERPINE1* expression, *PANX1* methylation, and *CDKN2A* deletion, we were able to identify subsets of patients with particularly poor outcomes, especially in the BRCA, GBMLGG, SARC, and STES cohorts. For instance, in breast cancer, patients whose tumors expressed high levels of *KIF20A* had a median survival time of 3430 days, while patients whose tumors also ‐harbored *CDKN2A* deletions, low levels of *PANX1* methylation, and had high levels of *SERPINE1* expression had a median survival time of 1318 days (Figure 6E). In total, these results demonstrate that combining proliferation-dependent and ‐independent features measured on different genomic platforms can greatly improve patient stratification.

## Discussion

In the developed world, cancer kills approximately one in five people (*39*). Yet, despite the dire threat that a cancer diagnosis often portends, recent evidence suggests that a growing number of patients are over-treated, and receive debilitating surgery or chemotherapy for ultimately benign conditions (*40*, *41*). Improved methods to discriminate between indolent and lethal tumors will protect quality-of-life and can help identify the subset of patients who would benefit most from aggressive treatment.

As cancers arise due to the accumulation of mutations in growth-promoting oncogenes and growth-inhibitory tumor suppressors, the presence and diversity of these mutations may be expected to dictate a tumor’s natural course. Surprisingly, our data suggest that they do not. For numerous oncogenes and tumor suppressors that play crucial roles in tumor development – including *KRAS, PIK3CA, CDKN2A*, and many more – the presence of mutations in these genes is uncorrelated with patient prognosis. However, our analysis includes several key caveats that underlie the potential benefits of continued sequencing efforts. First, oncogenic mutations can dictate susceptibility to targeted therapies, which are likely to play an increasingly-important role in cancer management (*26*). Secondly, our analysis of gliomas identified several potential instances of pathological misdiagnosis, with profound implications for patient survival. In certain tissues, mutational patterns can differentiate between cancer subtypes, and can therefore contribute to disease identification and staging. Finally, tumors themselves are composed of sub-clonal populations that harbor distinct sets of mutations, and recent evidence suggests that cancer heterogeneity can influence clinical course (*42*). Thus, interrogating the mutational spectrum at the sub-clonal level may identify prognostic mutations not distinguished in our population-based analysis. Nonetheless, our results suggest that with current technology, other molecular markers are likely to harbor more prognostic information than most mutations, even in potent driver genes.

Despite the limited stratification value of mutations in driver genes, copy number alterations of these same genes are broadly prognostic. Why would the amplification or deletion of a gene be associated with outcome, when a mutation in that same gene is not? We hypothesize that many of the driver gene mutations found in cancer are strictly required for tumorigenesis. In several tissues, a mutation to activate Ras/MAPK signaling – in *EGFR, KRAS, BRAF, PIK3CA*, *NF1*, or another related gene – seems to be necessary for tumor formation (*43*, *44*). In contrast, copy number changes in these genes are typically not required for tumorigenesis, but can instead modulate the degree of pathway activation or inactivation. In one well-studied example, a single *KRAS* mutation has been found to be sufficient to cause lung cancer, while additional copies of *KRAS* increase malignant growth by enhancing tumor metabolism (*45*). We speculate that this phenomenon may be common across driver genes, and copy number alterations at these loci are naturally selected to increase or decrease pathway output. Our results also imply that gene copy number – which can be imputed from common sequencing pipelines (*46*) – should be analyzed as a routine part of clinical sequencing efforts.

Careful analysis has shown that many prognostic gene signatures predominantly report mitotic activity in a tumor, and thus may provide little useful information not captured by Ki-67 staining or directly counting mitotic figures (*22*). Our findings underscore the broad importance of proliferation in risk assessment in most cancer types, though we note that some cohorts (LUSC, OV, and COADREAD, in particular) were poorly stratified by cell cycle markers. Prognosis in these cancers may depend on other factors, including the quality of surgical excision in ovarian cancer (*47*) or stromal signaling in colorectal cancer (*48*), that were not assessed in the TCGA. Nonetheless, our analysis demonstrates that the confounding effects of cell proliferation extend beyond protein-coding genes, and can influence the prognostic power of mutations, CNAs, methylated loci, and non-coding transcripts. Furthermore, we found that controlling for mitotic activity reveals a deeper layer of prognostic genes with roles in angiogenesis, cell motility, and invasion. These genes, rather than the “top” layer of prognostic cell cycle factors, may directly influence metastatic capacity, and warrant further study. Finally, several CNAs and methylated loci remained associated with survival in multivariate models when combined with a proliferation meta-gene. Thus, our analysis underscores the potential power to stratify patient risk by combining multiple complementary genome analysis platforms.

## Supplemental Figure Legends

**Figure S1.**
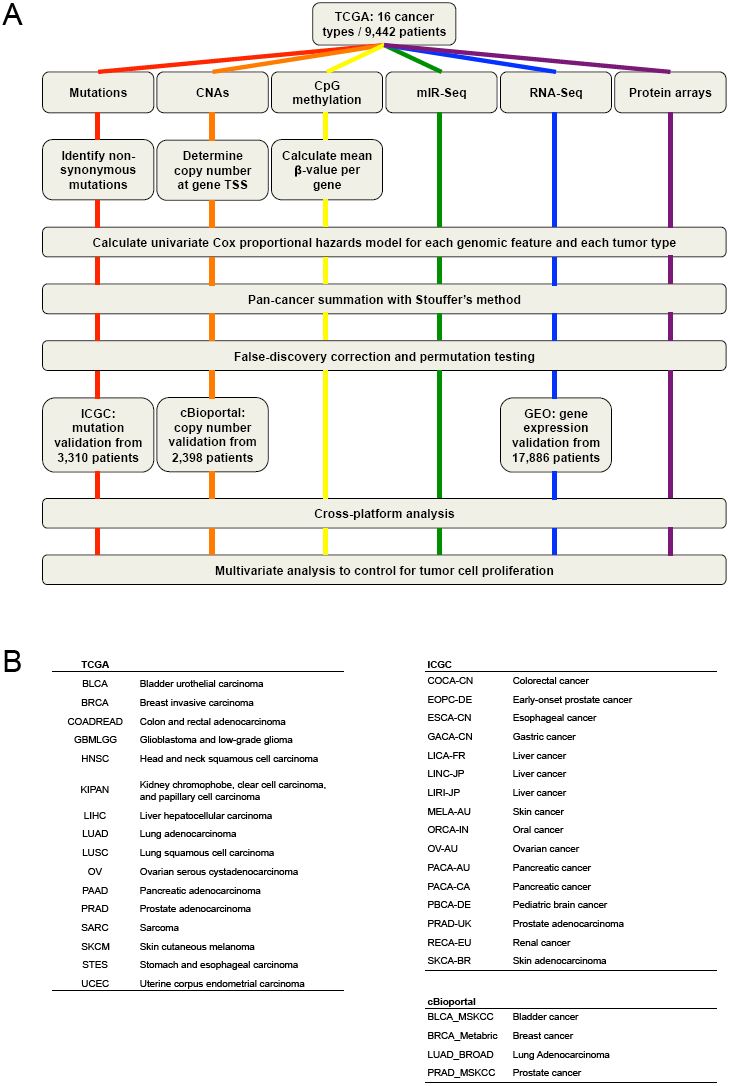
A schematic of the pan-cancer survival analysis pipeline and the datasets used. (A) An outline of the data processing and analysis performed in this report is presented. 16 tumor types from TCGA were used as a discovery cohort, while datasets from ICGC, cBioportal, and GEO were used as validation cohorts. (B) The cancer types and study abbreviations used in the TCGA, ICGC, and cBioportal datasets. More information on each cohort is included in Table S1.

**Figure S2.**
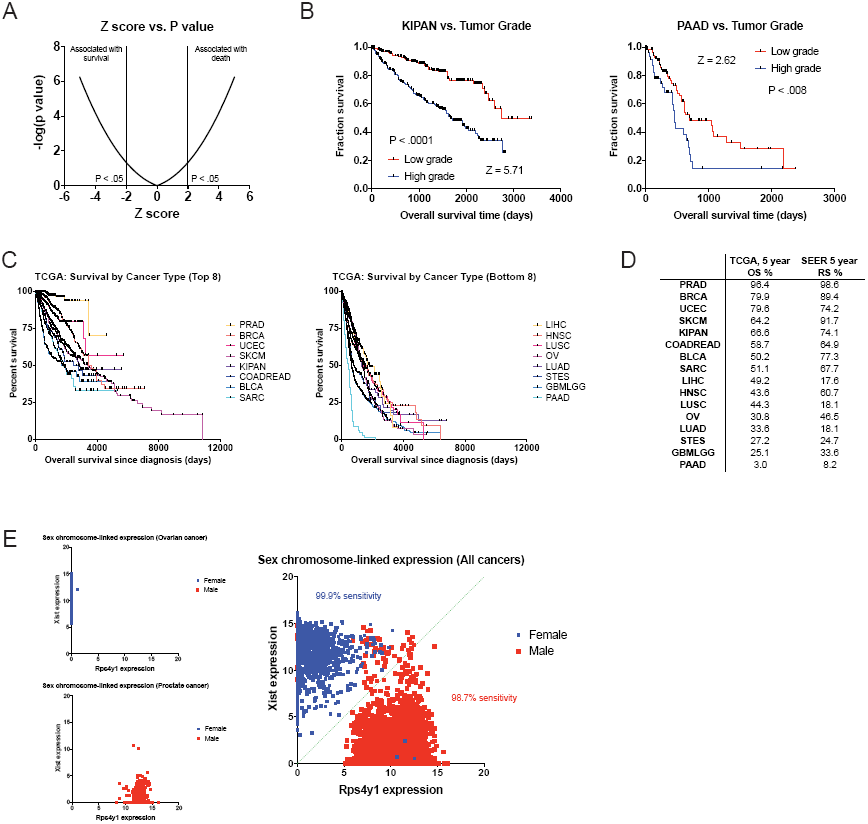
Tumor type-specific survival patterns and chromosome-specific gene expression patterns verify the accuracy of the TCGA clinical annotations. (A) The relation between Z score and P value for −5 ≤ Z ≤ 5 is displayed. (B) Kaplan-Meier plots showing a highly-significant survival difference (tumor grade in kidney cancer; Z = 5.71, P < .0001) and a moderately-significant survival difference (tumor grade in pancreas cancer; Z = 2.62, P < .008) are displayed. (C) Kaplan-Meier plots showing the survival time post-diagnosis for the eight cancers with the most favorable outcomes (left) and the eight cancer with the most dismal outcomes (right) in the TCGA dataset. (D) A comparison of the 5-year overall survival times from the TCGA with the 5-year relative survival times per cancer type from the Surveillance, Epidemiology, and End Results program (SEER) are displayed. (E) Scatter plots showing the expression of the X chromosome-encoded *XIST* transcript and the Y chromosome-encoded *RPS4Y1* transcript in ovarian cancer (OV), prostate cancer (PRAD), and across all 16 tumor types are displayed. As *Xist* is specifically expressed in cells that contain two or more copies of the X chromosome, this two gene combination has been shown to be effective as discriminating a patient’s chromosomal sex on the basis of gene expression (*13*, *28*). Accordingly, in our analysis, nearly all female patients have high *XIST* expression and low *RPS4Y1* expression, while nearly all male patients have high *RPS4Y1* expression and low *XIST* expression.

**Figure S3.**
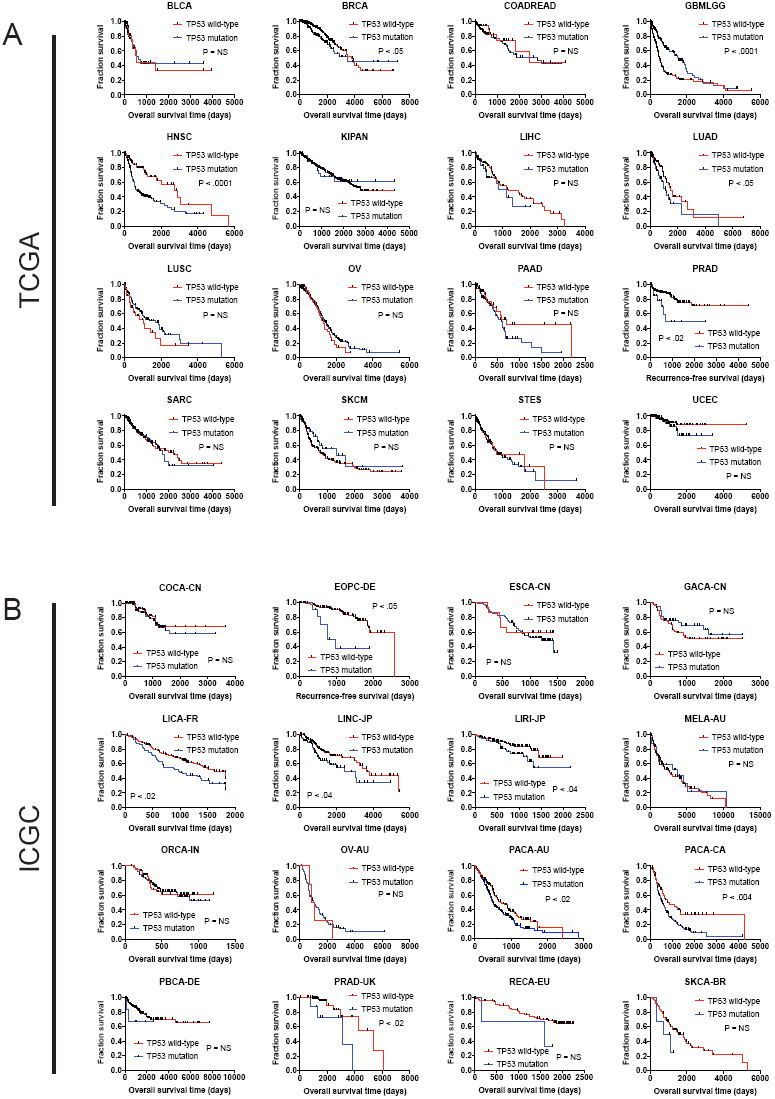
The mutation status of *TP53* is associated with outcome in multiple cancer types. (A) Kaplan-Meier plots of *TP53*-mutant and *TP53*-WT tumors in every cancer type from the TCGA set. *TP53* is associated with outcome in five of 16 cancer types (BRCA, GBMLGG, HNSC, LUAD, and PRAD). (B) Kaplan-Meier plots of *TP53*-mutant and *TP53*-WT tumors in every cancer type from the ICGC dataset. *TP53* is associated with outcome in seven of 16 cancer types (EOPC-DE, LICAFR, LINC-JP, LIRI-JP, PACA-AU, PACA-CA, and PRAD-UK).

**Figure S4.**
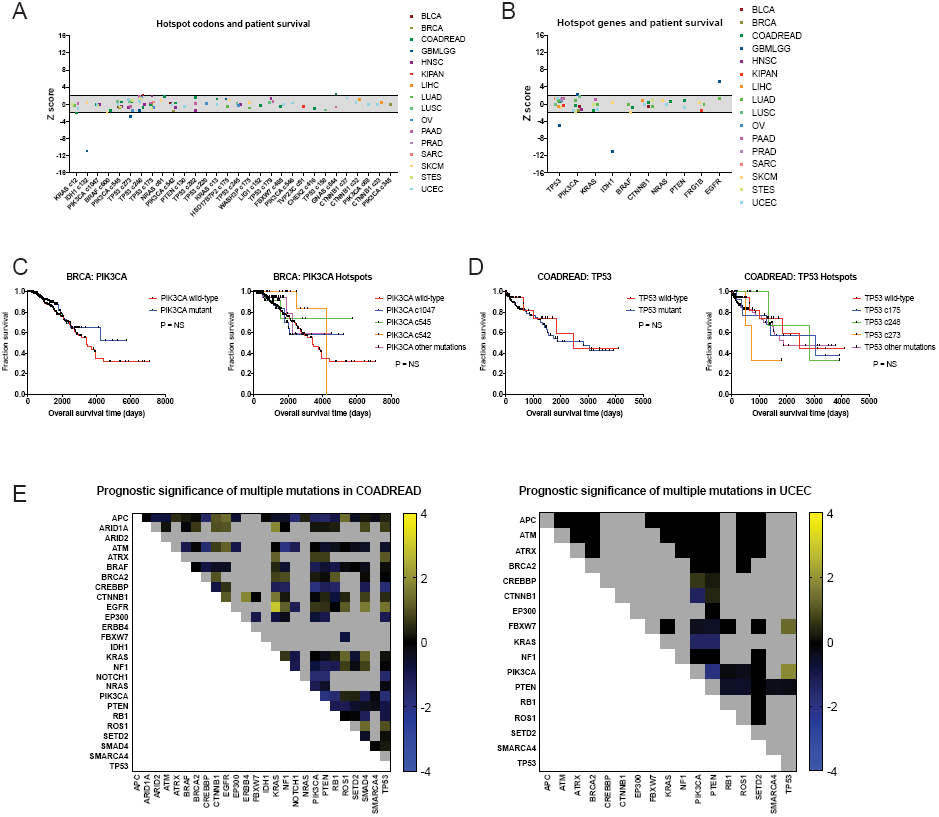
Hotspot mutations and mutations in multiple cancer driver genes are generally not associated with clinical prognosis. (A) Z scores of the 30 most frequently-mutated codons across 16 tumor types from the TCGA are displayed. Z scores were calculated comparing survival times between patients harboring a mutation at that codon and patients that did not have a mutation at that codon for all cancer types in which a particular codon was mutated in ≥2% of samples. Dotted lines are plotted at Z = ±1.96, corresponding to P < .05. (B) All codons mutated in ≥5 patients (“hotspot” mutations) were pooled together, and the ten genes harboring the most “hotspot” mutations were determined. Z scores were calculated comparing patients harboring any “hotspot” mutation in a specified gene and patients that do not have any “hotspot” mutation in that gene. Z scores are plotted for the 16 tumor types in the TCGA dataset. Dotted lines are plotted at Z = ±1.96, corresponding to P < .05. (C) Kaplan-Meier curves are plotted comparing BRCA patients with any mutation in *PIK3CA* vs. those with no mutations in *PIK3CA (*left), and BRCA patients with various “hotspot” mutations in *PIK3CA* vs. those with no “hotspot” mutations in *PIK3CA (*right). Neither method of dividing patients identifies a subgroup with a significantly different clinical outcome. (D) Kaplan-Meier curves are plotted comparing COADREAD patients with any mutation in *TP53* vs. those with no mutations in *TP53 (*left), and COADREAD patients with various “hotspot” mutations in *TP53* vs. those with no “hotspot” mutations in *TP53 (*right). Neither method of dividing patients identifies a subgroup with a significantly different clinical outcome. (E) Heatmaps of double-mutation combinations in COADREAD and UCEC are plotted. Among 30 frequently-mutated cancer driver genes, 26 are mutated in ≥2% of COADREAD patients and are not individually correlated with prognosis, while 17 are mutated in ≥2% of UCEC patients and not individually correlated with prognosis. Z scores were calculated by comparing survival times of patients harboring a non-silent mutation in two different cancer driver genes vs. patients that did not harbor a non-silent mutation in those two genes. The blue and yellow color bars indicate Z scores.

**Figure S5.**
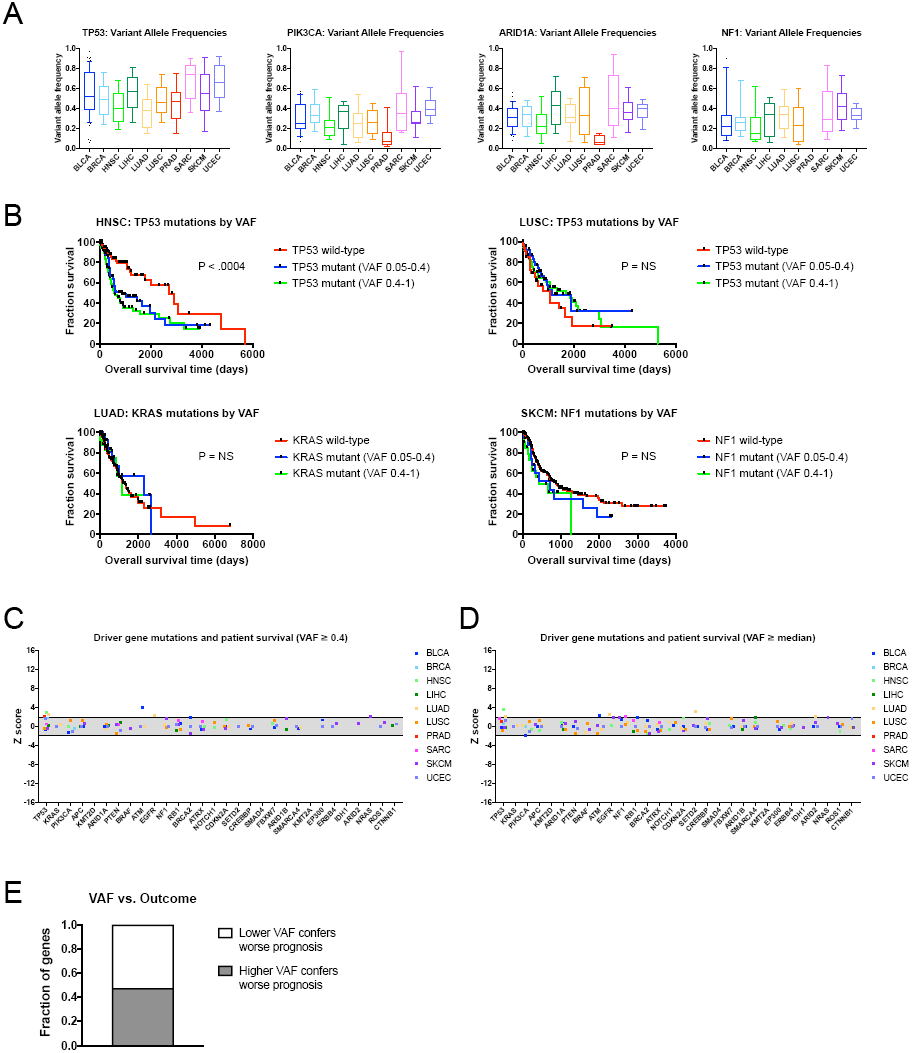
Mutations with high variant allele frequencies are no more prognostic than mutations with low variant allele frequencies. (A) The variant allele frequencies were calculated for all genes in 10 TCGA cohorts. Box plots of the VAFs of four common cancer drivers (TP53, PIK3CA, ARID1A, and NF1) are displayed. Boxes represent the second and third VAF quartiles, while error bars indicate the 10^th^ and 90^th^ percentiles. (B) Kaplan-Meier curves of patient outcomes versus mutation VAF. TP53 mutations are associated with outcome in HNSC, and this relationship is true for both high-prevalence (VAF ≥0.4) and low-prevalence (VAF < 0.4) mutations (top left plot). Similarly, no significant prognostic information is conferred by TP53 mutations in LUSC, KRAS mutations in LUAD, or NF1 mutations in SKCM, regardless of the prevalence of the mutation. (C) Z scores of the 30 most frequently-mutated cancer driver genes are displayed. Z scores were calculated by regressing survival times between patients harboring wild-type and mutant copies of a gene if a gene was mutated in ≥2% of samples per tumor type. However, a gene was only considered “mutated” for this analysis if its VAF was ≥0.4. Dotted lines are plotted at Z = ±1.96, corresponding to P < .05. (D) Z scores of the 30 most frequently-mutated cancer driver genes are displayed. Z scores were calculated by regressing survival times between patients harboring wild-type and mutant copies of a gene if a gene was mutated in ≥4% of samples per tumor type. For this Z score analysis, only mutations with VAFs greater than or equal to the median VAF for a particular gene in a particular cancer type were included. Dotted lines are plotted at Z = ±1.96, corresponding to P < .05. (E) We compared the Z scores that resulted from regressing only above-median VAFs and the Z scores that resulted from regressing only below-median VAFs, for the 30 cancer driver genes. In 62 instances, higher Z scores resulted from only considering above-median VAFs, while in 68 instances, higher Z scores resulted from only considering below-median VAFs.

**Figure S6.**
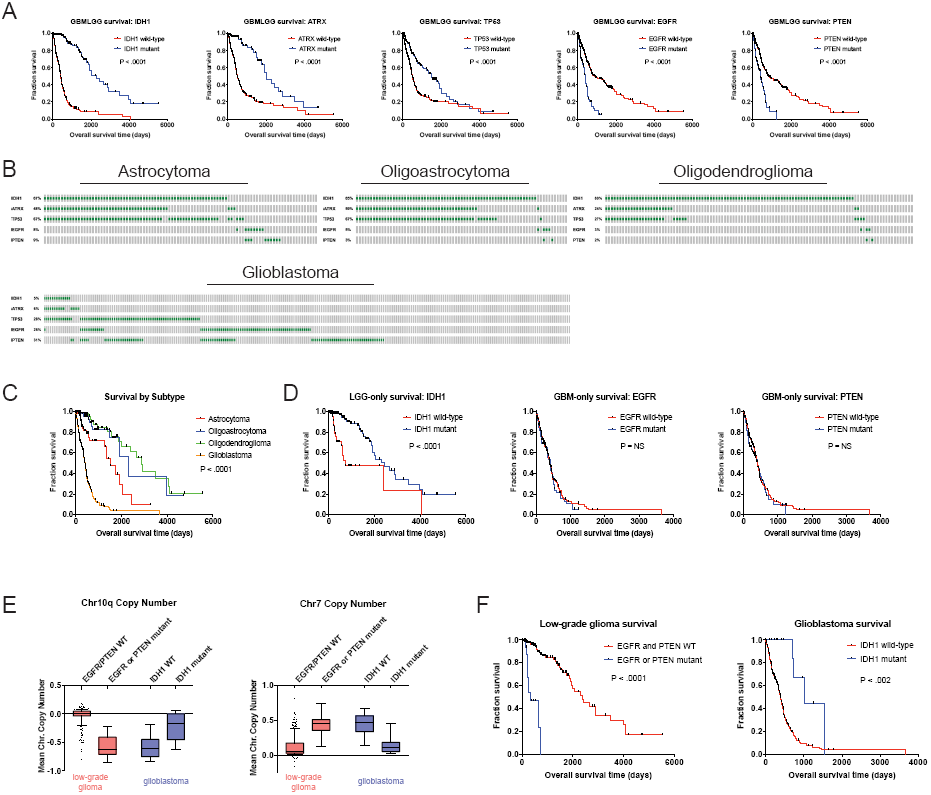
Subtype-specific mutation patterns in glioma correlate with survival time and can inform clinical diagnosis. (A) Kaplan-Meier curves of the five genes with the strongest survival associations in GBMLGG (*ATRX*, *EGFR*, *IDH1*, *TP53*, and *PTEN*) are displayed. (B) Mutation patterns according to glioma subtype are displayed. Gray boxes correspond to patients with a particular glioma subtype, while green dashes indicate the presence of a mutant copy of the specified gene. (C) Kaplan-Meier curves of patient survival according to glioma subtype are displayed.Glioblastoma confers the worst prognosis, while low-grade gliomas confer a more favorable prognosis. (D) Subtype-specific survival patterns partially explain the association between mutations and clinical outcome in glioma. When only considering patients with glioblastoma, patients with mutations in *EGFR* or *PTEN* have similar prognoses compared to patients with wild-type copies of these genes. However, among patients with low-grade glioma, patients with wild-type *IDH1* exhibit worse overall survival compared to patients with *IDH1* mutations. (E) Patients with subtype-discordant mutation patterns also exhibit chromosome copy number changes characteristic of a different glioma subtype. Chromosome 10q is frequently lost in glioblastoma (*33*). Patients diagnosed with low-grade glioma who harbor *PTEN* or *EGFR* mutations tend to also lose Chr10q, while patients diagnosed with glioblastoma who harbor mutations in *IDH1* tend to have wild-type levels of Chr10q. In contrast, chromosome 7 is frequently gained in glioblastoma, and patients with subtype-discordant mutation patterns have Chr7 copy numbers that correspond to their mutational status, rather than their clinical diagnosis. (F) Kaplan-Meier curves of patients diagnosed with low-grade glioma who harbor *PTEN* or *EGFR* mutations (left) and of patients diagnosed with glioblastoma who harbor *IDH1* mutations (right). Survival curves mimic patients’ mutational status rather than their clinical diagnosis.

**Figure S7.**
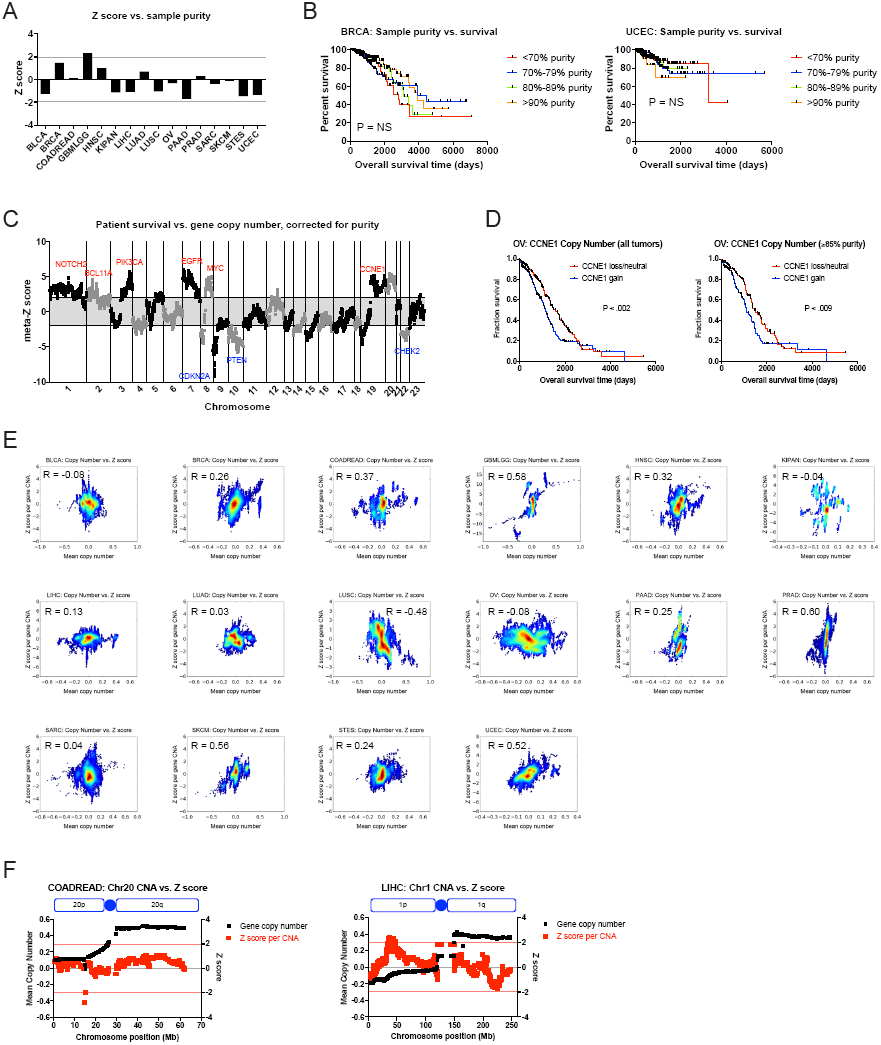
Figure S7. The prognostic value of cancer CNAs is independent of tumor sample purity. (A) A bar graph showing Z scores obtained by regressing sample purity, as measured by IHC, against patient survival. Dotted lines indicate Z scores of 1.96 and −1.96, corresponding to a P value < .05. (B) Representative Kaplan-Meier plots of patient survival in BRCA and UCEC, split based on the purity of the sample analyzed in the TCGA analysis. (C) Z scores from multivariate models including copy number and tumor purity from the 16 cancer types from the TCGA were combined using Stouffer’s method, and then the resulting meta-Z scores were plotted against the chromosomal location. Genes were binned by average Z score into groups of 5 for visualization. (D) CNAs remain prognostic even in pure tumor samples. The survival of OV patients according to *CCNE1* copy number is plotted, using data from either all patients (left) or only patients with tumor samples with ≥85% purity. (E) For each of the 16 TCGA cancer cohorts profiled, the mean copy number per gene is plotted against the Z score regressing copy number against outcome. The absence of a strong correlation between copy number and outcome indicates that only certain CNAs drive patient prognosis. (F) Representative comparisons of mean copy number vs. Z score. Shown here are chromosome 20 from COADREAD and chromosome 1 from LIHC. Mean copy number per gene is plotted in black, while the Z score at each gene is plotted in red. In these patient cohorts, 1q and 20q are commonly gained but are not associated with grim prognosis.

**Figure S8.**
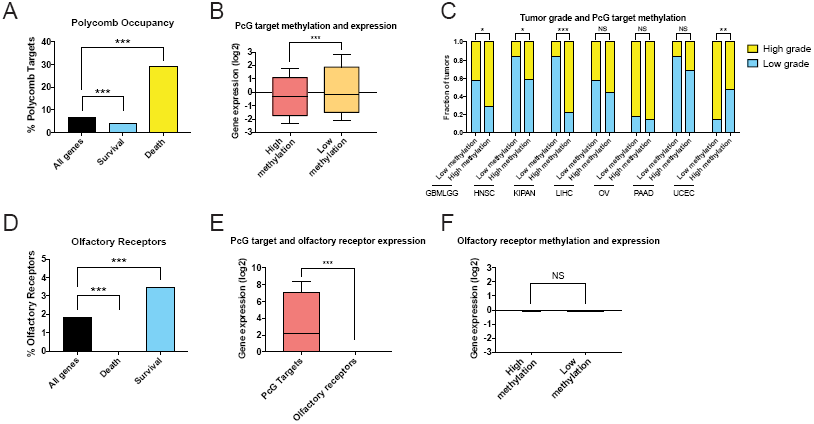
Methylation of Polycomb targets is associated with high tumor grade and patient mortality. (A) Death-associated methylation sites are enriched for genes bound by Polycomb-component Suz12 (*36*) in hES cells. ***, P < .0005 (hypergeometric test). (B) Death-associated Polycomb target genes were split based on the average level of methylation at that gene within each tumor types. Across all death-associated Polycomb targets, lower levels of DNA methylation were associated with higher levels of gene expression.Boxes represent the second and third expression quartiles, while error bars indicate the 10^th^ and 90^th^ percentiles. ***, P < .0005 (Student’s t-test). (C) Each tumor type was split into two groups based on the average level of methylation of all death-associated Polycomb targets. High-grade (g3 or g4) GBMLGG, HNSC, and KIPAN cancers were enriched for tumors with high levels of Polycomb target methylation. *, P < .05; **, P < .005; ***, P < .0005 (Fischer’s exact test). (D) Survival-associated methylation sites are enriched for genes encoding proteins with olfactory receptor activity. ***, P < .0005 (hypergeometric test). (E) A box-and-whiskers plot of the log2-transformed RSEM expression values of olfactory receptor genes and Polycomb targets across the TCGA datasets. Boxes represent the second and third expression quartiles, while error bars indicate the 10^th^ and 90^th^ percentiles. (F) Genes encoding survival-associated olfactory receptors were split based on the average level of methylation at that gene within each tumor types. The level of DNA methylation across all survival-associated olfactory receptors was not significantly correlated with gene expression at that locus.

**Figure S9.**
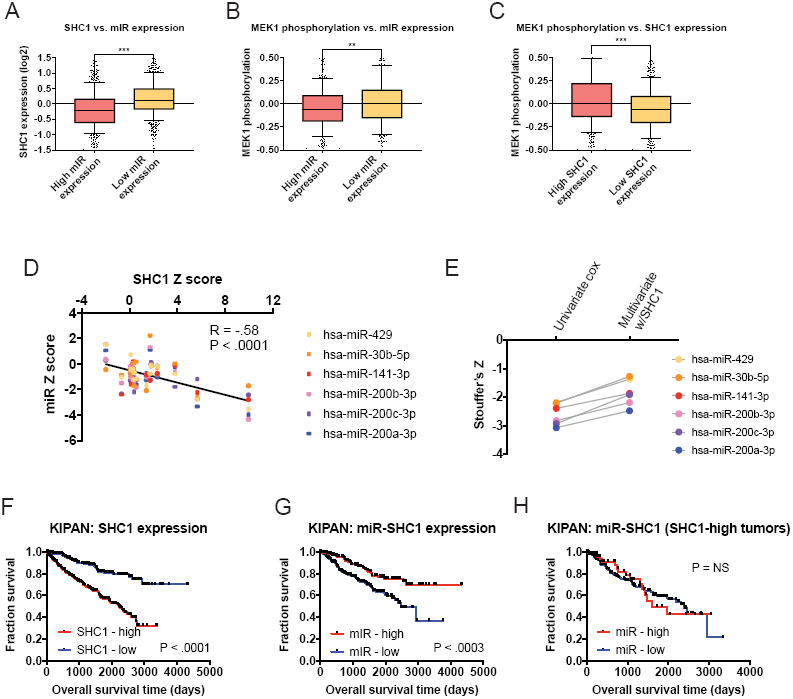
microRNAs targeting SHC1 are associated with favorable prognosis. (A) Tumors with high expression of the SHC1-targetting microRNAs hsa-miR-429, hsa-miR-30b-5p, hsa-miR-141-3p, hsa-miR-200b-3p, hsa-miR-200c-3p, and hsa-miR-200a-3p (henceforth, mIR-SHC1) express lower levels of *SHC1*. Boxes represent the second and third expression quartiles, while error bars indicate the 10^th^ and 90^th^ percentiles. ***, P < .0005 (Student’s t-test). (B) Tumors with high expression of mIR-SHC1 have lower levels of MEK1 phosphorylation, as judged by RPPAs. **, P < .005 (Student’s t-test). (C) Tumors with lower levels of *SHC1* expression have lower levels of MEK1 phosphorylation, as judged by RPPAs. ***, P < .0005 (Student’s t-test). (D) Z scores for each mIR-SHC1 in each of the 16 TCGA cohorts were plotted against Z scores for *SHC1* expression. (E) Multivariate models were constructed containing *SHC1* expression and each individual mIR-SHC1 in each of the 16 TCGA cohorts. Pan-cancer Stouffer’s Z scores were calculated across cancer types, and univariate vs. multivariate Z scores are plotted. (F) In the pan-kidney TCGA cohort, high expression of *SHC1* is associated with shorter overall survival. (G) In the pan-kidney TCGA cohort, high expression of mIR-SHC1 is associated with longer overall survival. (H) In the pan-kidney cohort, among only those patients in the top half of *SHC1* expression, assessing mIR-SHC1 expression is no longer associated with outcome, suggesting that *SHC1* and mIR-SHC1 capture largely the same prognostic information.

**Figure S10.**
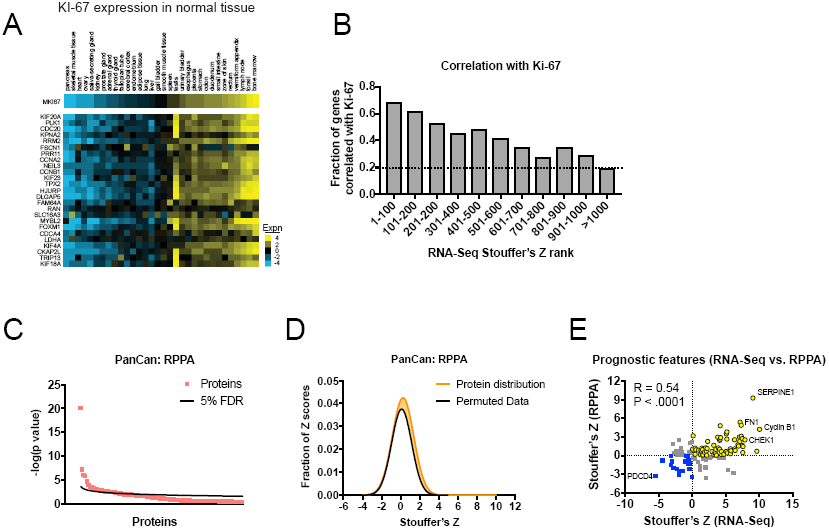
Prognostic transcripts are correlated with Ki-67 expression in normal tissue and are commonly associated with outcome at the protein level. (A) A heatmap of the expression of the 25 top-scoring transcripts in normal tissue (*49*) sorted according to *MKI67* expression. (B) A bar graph of the fraction of genes correlated with *MKI67* expression in normal tissue, according to the rank of the gene in the RNA-Seq pan-cancer analysis. The dotted line at Y = 0.2 indicates the fraction of all genes significantly correlated with *MKI67*. (C) Univariate Cox models were calculated for every protein or protein modification measured in the TCGA. The significance of the top-scoring proteins are plotted against a 5% false-discovery threshold calculated using the Benjamini-Hochberg method. The complete list of Z scores is presented in Table S2F. (D) A histogram of meta-Z scores for proteins are plotted against a histogram of meta-Z scores from ∽1,000,000 random permutations of the underlying patient data. (E) A scatter plot of meta-Z scores for individual genes, calculated using either RNA-Seq or RPPA values. Genes were only included if the antibody used in RPPA analysis recognized a specific protein without any post-translational modifications (cleavage, phosphorylation, etc.).

**Figure S11.**
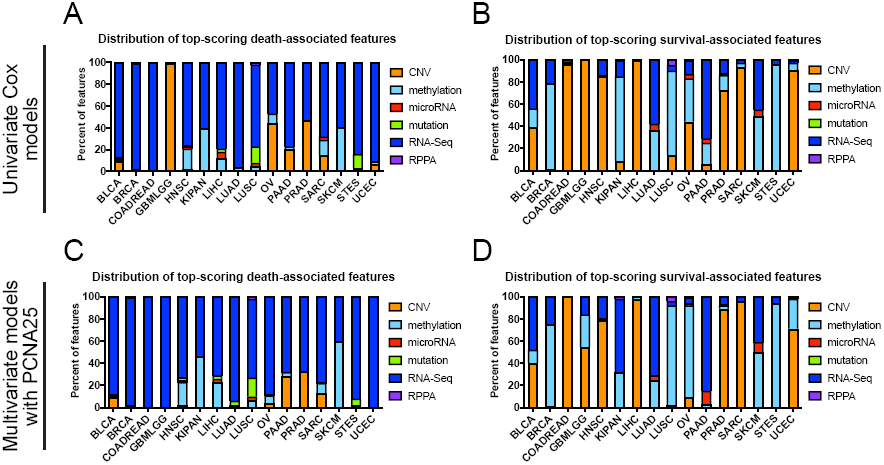
Prognostic features in individual tumor types. (A and B) A bar graph displays the distribution of feature types among the 100 genomic features that exhibited the strongest association with (A) death from cancer or (B) cancer survival in univariate Cox models. (C and D) A bar graph displays the distribution of feature types among the 100 genomic features that exhibited the strongest association with (A) death from cancer or (B) cancer survival in multivariate Cox models with the PCNA25 meta-gene.

**Figure S12.**
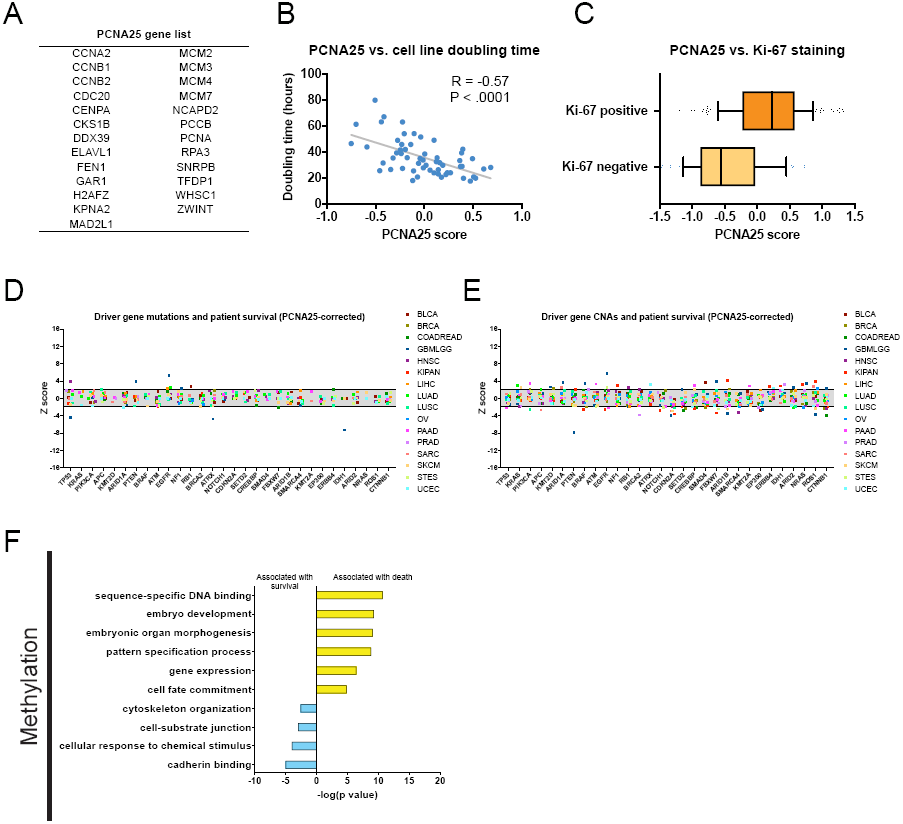
Multivariate analysis with a proliferation meta-gene. (A) A list of the 25 genes that comprise the PCNA25 gene signature (*38*). (B) The PCNA25 gene score correlates with the doubling time of the NCI-60 cell panel in culture (*50*). (C) The PCNA25 gene score correlates with Ki-67 staining via IHC (from breast cancer dataset GSE21653) (*51*). (D) Z scores from multivariate models with PCNA25 of 30 frequently-mutated cancer driver genes in 16 tumor types from the TCGA are displayed. Z scores were calculated comparing survival times between patients harboring wild-type and mutant copies of a gene for all genes mutated in ≥2% of samples per tumor type. Dotted lines are plotted at Z = ±1.96, corresponding to P < .05. The complete list of Z scores is presented in Table S5A. (E) Z scores from multivariate models with PCNA25 regressing gene copy number against patient outcome for the 30 most frequently-mutated cancer driver genes are displayed. Dotted lines are plotted at Z = ±1.96, corresponding to P < .05. The complete list of Z scores is presented in Table S5B. (F) Gene ontology terms over-represented among methylation sites that are significantly associated with either patient survival or patient death, after PCNA25 correction, are displayed.The complete list of Z scores is presented in Table S5C.

## Materials and methods

### Data Sources

Patient cohorts analyzed in this study are listed in Table S1. For the TCGA analysis, pre-processed files from the Broad Institute TCGA Firehose were used (*52*). For the TCGA genomic copy number analysis, we used the HG19 segmented SCNAs, corrected for germline SCNAs. For methylation, we used the average β-value per gene. For microRNAs, we used log2-normalized pre-processed mature microRNA annotations. For RNA-Seq, we used RNASeq-v2 RSEM data, normalized by gene. Overall survival was used as a clinical endpoint for all cancer types except PRAD; due to the few deaths in the PRAD cohort, “days to biochemical recurrence” was used as an endpoint. For all cancers, survival or follow-up time from diagnosis were corrected for the days to sample procurement. Only primary tumors (indicated with a “01” in the patient barcode) were used for every cancer type except SKCM; for this cancer, few primary samples were available, so metastatic samples (indicated with a “06” in the barcode) were included for patients in which no primary tumor was available. “Percent_tumor_cells” was used to infer sample purity.

Mutation and clinical data from Release 25 of the International Genome Consortium were downloaded from the ICGC Data Portal (*53*). Overall survival was used as a clinical endpoint for all cohorts except EOPC-DE; due to the few deaths in this cohort, recurrence-free survival was used as an endpoint. Cohorts were chosen based on the availability of WGS or WES data, and were included if they came from a cancer type comparable to the types that were studied in our TCGA analysis. Copy number and clinical data from cBioportal were downloaded from www.cbioportal.org (*54*).

Data from the Gene Expression Omnibus were downloaded from https://www.ncbi.nlm.nih.gov/geo (*55*). For cohorts with multiple clinical endpoints, we used 1) Overall survival, 2) Disease-specific survival, or 3) Recurrence or metastasis-free survival, based on availability. Probeset definitions were downloaded from GEO.

### Overall Analysis Strategy

All processing and analysis was performed using Python. Cox proportional hazard analysis used the R survival package (https://cran.rproject.org/web/packages/survival/index.html) to compute Z scores and p values. The rpy2 project was used to control R from python, allowing seamless integration of Z score calculations with data processing and pan-cancer analysis. Pandas DataFrames were used as the primary structure for storing and manipulating data. Additionally, native numpy methods and arrays were for used occasionally for efficiently storing strictly numerical data, for example, as input to Cox proportional hazards models. The statsmodels package (statsmodels.org) was used for false discovery correction using the Benjamini-Hochberg procedure. Microsoft Excel was occasionally used for final data processing and examination, so a single apostrophe was added before gene names in intermediate data processing steps to protect genes from auto-formatting (*56*).

Code was structured to allow ease of internal reuse and reproducibility of results. Cox univariate proportional hazards, Cox multivariate proportional hazards, Kaplan-Meier, and Stouffers analysis methods were factored into an analysis library, taking as input the data required to perform the computation as numpy arrays or pandas DataFrames.

### TCGA Analysis

In addition to the code for statistical analyses, code for processing TCGA clinical files was factored into a common library. This approach allowed the same TCGA clinical file processing code to be executed across a variety of platform analyses, ensuring identical behavior for each platform. The TCGA clinical processing code selected the relevant clinical endpoints and sample procurement data. The processing translated the available clinical data into the required format for Cox proportional hazard models: an endpoint and a censor value for each patient. Code to select tumor samples based on cancer type was also included in this library. For every dataset analyzed except skin cutaneous melanoma (SKCM), only primary tumor samples were included. In SKCM, metastases samples were also included if there was no primary tumor sample data available for a patient.

Four platform analyses were nearly identical, and thus common code was written to perform these: RPPA, RNA-Seq, methylation, and microRNA analyses were performed using a single codebase with slightly varying input parameters. To accomplish this generalization, the code has some limited dependencies on the file structure of the raw data. For example, the directory names of each platform must conform to a standard in defined in the code, and the individual clinical and platform files must have names that predictably include the cancer type. For all four of these platforms, Cox proportional hazards were only calculated if the gene in question appeared at least 10 times in a cancer type.

The remaining two TCGA platforms (mutation and copy number) needed to be treated independently. Raw input data for the mutation analysis needed additional preprocessing before Cox proportional hazard models could be calculated. This preprocessing included removing per-patient headers throughout the data and some data transposition. For all analyses using TCGA mutation data, mutations annotated as silent were excluded. Genes were only included in downstream analyses if they were mutated in 2% or more patients in a cancer type.

Raw input data for copy number analysis also required substantial preprocessing. Copy number input data consists of per-patient, per-chromosome location maps of copy numbers (hg19 downloaded from the UCSC Genome Browser (*57*)). These maps were converted to a single copy number value for each gene. We created an interval tree (using the intervaltree python package, https://pypi.python.org/pypi/intervaltree) of the location maps for each chromosome and used the appropriate HGNC to convert chromosome locations to genes for each patient. We used the gene’s transcriptional start site position to look up in the interval tree the copy number value for a gene. This analysis produced an intermediate file of a similar form to the other TGCA platforms, which allowed for straightforward pan-platform analysis. For copy number analysis, Cox proportional hazards were only calculated for genes that appeared at least 10 times in a cancer type.

### TCGA Meta-gene Analysis

In addition to performing univariate Cox models described above, we performed meta-gene analysis on each of the six platforms for PCNA25, *SHC1*, *KIF20A*, and *MKI67*. Each meta-gene for multivariate Cox analysis was preprocessed to give a single combined value for each patient. Normalization required to calculate meta-gene values was performed by taking the log2 transform of each gene’s expression, clipping at 0 and subtracting the mean of each gene across patients within each cancer type. We then averaged the normalized expression values of each gene in the meta-gene to produce a single meta-gene value for each patient. This value was used as the second variable in Cox multivariate proportional hazard models for each of the six platforms.

### Pan-cancer TCGA Analysis

For each platform and analysis type, we performed a pan-cancer analysis. This analysis created a single Z score for each gene by combining the per gene Z scores from each cancer type using Stouffer’s method. To perform Stouffer’s method, we took the sum of the Z scores for a single gene and divided that sum by the square root of the number of cancer types with Z scores for the gene. This meta-Z score was then compared against meta-Z scores obtained similarly from other platform analyses.

### Pan-platform TCGA Analysis

Pan-platform (RNA-Seq, methylation, RPPA, microRNA, copy number, mutation) TCGA analyses primarily used the raw data from each platform. For copy number data, we instead we used the preprocessed data described above. These pan-platform analyses required distinct normalizations of the data per-platform to provide comparable values. For copy number, RPPA, and methylation values, we normalized to 0 by subtracting the mean value of each gene within a particular cancer type. For microRNAs, we clipped values at 0 and subtracted the mean of each gene across patients. RNA-Seq data was normalized by taking the log2 transform, clipping at 0 and subtracting the mean of each gene across patients.

### Additional TCGA Mutation Analyses

Beyond the standard univariate and multivariate analyses performed for the other platforms, we performed three additional analyses on mutation data: double mutation combination Z scores, hotspot codon Z scores, and Z scores corrected for VAFs.

For double mutation Z scores, we took the top 30 most common cancer driver genes and pairwise combined them in every possible way. We then calculated Cox proportional hazards for each pair of genes, where a patient was considered to have a pairwise mutation if and only if both genes were non-silently mutated for that patient. Z scores were only calculated for a pair if (1) neither gene in the pair was statistically significant alone in the univariate analysis and (2) if both genes were mutated together in at least 10 patients.

Per codon Z scores were calculated for a selected set of hotspot codons. Most cancer types were available in HG37, so HG37 mutation positions were used to locate codons. Mutations for OV and COADREAD were only available in HG36, so gene positions were converted to HG37 before codon processing. Per codon Z scores were calculated by first identifying patients with mutations in the relevant gene, then selecting from that set of patients those whose mutations were in the codon of interest. If 2% of patients or more had mutations in the selected codon, a Z score was calculated.

VAFs were calculated for 10 of the TCGA cancer types. We analyzed VAF data in two ways. First, we calculated Z scores, only counting a gene as mutated if its VAF was greater than or equal to 0.4. Secondly, we identified the median VAF score per gene, and calculated Z scores only counting a gene as mutated if its VAF was equal to or greater than the median VAF for that gene.

### CBioPortal Analysis

CBioPortal was structured similarly to the TCGA analyses, though data processing was not factored into an independent library since each of these datasets was only used in one analysis. One CBioPortal cancer type, blca_mskcc, required initial preprocessing in the manner described above for TCGA copy numbers. CBioportal copy number Z scores were calculated only when a gene appeared at least 10 times in a cancer type.

### ICGC Analysis

ICGC analysis was structured similarly to CBioPortal analysis. Mutations were only included in downstream analyses if they were annotated as one of these types: disruptive inframe deletion, disruptive inframe insertion, frameshift variant, inframe deletion, missense variant, splice acceptor variant, splice donor variant, stop gained, or stop lost. Z scores were calculated when at least 2% of patients in a cancer type were mutated.

### GEO analysis

Survival-associated microarray datasets were found by searching the Gene Expression Omnibus. (*55*) Processed values were downloaded and log2-normalized when appropriate.GEO Z scores were calculated for genes where at least 50% of patients had non-missing expression data. Z scores from probes corresponding to the same gene were combined by averaging. Pan-cancer analysis of GEO data was performed using Stouffer’s method (as described above) to combine Z scores across cohorts for a single cancer type, then again across cancer types to create a single pan-cancer GEO Z score for each gene.

### Code

Code is available on github at https://github.com/joan-smith/genomic-features-survival, and supplemented by https://github.com/joan-smith/microarray-proportional-hazards for GEO Cox proportional hazard calculation.

### Additional data sources and tools

Frequently-mutated cancer driver genes were acquired from Zehir et al., 2017 (*29*).Suz12-bound sites were acquired from Lee et al., 2006 (*36*). Gene expression from normal human tissue was acquired from dataset E-MTAB-2836 (*49*) from the RNA-Seq Atlas (*58*). The PCNA25 gene list was downloaded from Sheltzer, 2013 (*38*). Heatmaps were generated in Java Treeview (*59*). Density plots were performed with Python scripts using matplotlib (https://matplotlib.org). Gene Ontology analysis was performed using GProfiler (*60*) with a Benjamini-Hochberg correction, against a background list of all genes present in the particular assay. GSEA on MicroRNA targets was performed using miEAA with a minimum threshold of 7 microRNAs (*61*). Single base-pair mutations were mapped to codons using PolyPhen-2 (*62*).

